# The Leader of the Capsid protein from *Feline calicivirus* must be palmitoylated and form oligomers through disulfide bonds for efficient viral replication

**DOI:** 10.1101/2024.06.07.597891

**Authors:** Yoatzin Peñaflor-Téllez, Jaury Gómez de la Madrid, Erick Ignacio Monge-Celestino, Carolina Pérez-Ibáñez, Carlos Emilio Miguel-Rodríguez, Ana Lorena Gutiérrez-Escolano

**Affiliations:** Departamento de Infectómica y Patogénesis Molecular, Centro de Investigación y de Estudios Avanzados del IPN

**Keywords:** Feline calicivirus, LC protein, viroporin, palmitoylation, PDIA3, oligomers

## Abstract

*Feline calicivirus* (FCV), a member of the *Vesivirus* genus and a model for studying of the members of the *Caliciviridae*, has been used to understand the biology and viral replication of etiological agents of medical and veterinary importance. The leader of the capsid (LC) protein is exclusive to the *Vesivirus* genus members whose importance for a successful viral replication has been demonstrated; however, its cell localization, post-translational modifications, and specific functions throughout the viral replication cycle are poorly understood. We have determined that the expression of the LC protein from FCV in a virus-free system is located at the mitochondria and induces apoptosis; furthermore, the *in vitro* characterization of a purified LC protein showed that it has viroporin characteristics. Here, using an LC-reactive serum, the LC protein expression kinetics, cellular localization, post-translational modifications, and association with cellular PDI proteins in FCV infected cells was determined. We found that the LC protein is present on the membrane of infected cells and in the supernatants, independently of cell lysis, which suggests that it is actively secreted. Moreover, we found that LC protein is palmitoylated during infection and that palmitoylation inhibition alters its levels and subcellular localization. Finally, we determined that the LC protein forms oligomers dependent on disulfide bonds mediated by PDI activity. In addition, it interacts with PDIA3, a disulfide isomerase known as an important factor in the replication of several viruses. The results indicate that the LC protein from FCV might have multiple roles during FCV replication.

**IMPORTANCE:** Feline calicivirus (FCV) is a highly transmissible virus that represents a significant cause of upper respiratory infection in domestic and wild cats worldwide. FCV also serves as one of the most valuable models for studying calicivirus biology, as unlike most members of the family, known to cause diseases in animals and humans, it can be easily grown in cell culture. Since there are no efficacious vaccines or antivirals against most caliciviruses, studying their molecular biology and the relationship between viral and cellular components is essential for developing strategies for their prevention and control.

## Introduction

*Feline calicivirus* (FCV), a member of the *Vesivirus* genus in the *Caliciviridae* family, efficiently replicates in cell culture and has multiple reverse genetic systems. For the last 70 years, it has been one of the most important models for understanding the replication strategies of the members of the *Caliciviridae* family in infected cells (Peñaflor-Téllez et al., 2020). FCV is an etiological agent of upper respiratory tract pathologies in domestic and wild cats (Hofmann-Lehmann et al., 2022). In some cases, FCV infection can result in highly lethal systemic pathology that can easily spread in shelters or veterinary clinics (Bordicchia et al., 2021; Deschamps et al., 2015; Hurley et al., 2004; Pedersen et al., 2000). To this date, some vaccines confer some degree of protection against FCV pathology (Bergmann et al., 2019; Huang et al., 2010; Lesbros et al., 2013), but there is no commercial, effective treatment for ongoing infections, making it an important veterinary and wildlife concern.

Caliciviruses are non-enveloped RNA viruses that have positive-sense, single-stranded genomes that encode six non-structural (NS1-NS7) and two structural (VP1 and VP2) proteins. Although all viral proteins are encoded in the genomic RNA (gRNA), the structural proteins are translated late in infection from a subgenomic RNA (sgRNA) (Herbert et al., 1996). One marked difference between vesiviruses such as FCV and other members of the *Caliciviridae* family is the presence of a unique protein named the leader of the capsid (LC), which is translated from the sgRNA as an LC-VP1 precursor. This precursor is further processed by the viral protease-polymerase NS6/7 to generate both the mature LC and the major capsid protein VP1 (Carter et al., 1992; Sosnovtsev et al., 1998a). Although the LC does not form part of the mature FCV virions (Sosnovtsev & Green, 2000), it plays a significant role during infection.

In 2013, Abente et al. described that the LC protein is crucial for establishing the cytopathic effect during FCV infection and overall virus replication (Abente et al., 2013). The same research group found that FCV LC protein interacts with Annexin A2, a protein which our workgroup described as an important cellular factor for efficient replication (Santos-Valencia et al., 2019). We later determined that its ectopic expression results in its localization to the mitochondria and apoptosis triggering through the intrinsic pathway (Barrera-Vázquez et al., 2019). We performed different approaches to better characterize this protein and found that the LC protein is intrinsically toxic and forms disulfide-bond dependent homo-oligomers (Peñaflor-Téllez et al., 2022). This led us to suggest that the LC protein from FCV is a viroporin; however, more information regarding its expression, localization, and interaction with cellular components is needed to fully understand its role(s) in the FCV replication cycle.

In this work, we described the expression kinetics of the LC protein during infection. Moreover, we found that its subcellular localization in infected cells drastically differs from the LC protein-expressing cells. The LC protein was also found on the outer face of the cytoplasmic membrane and in the extracellular medium, suggesting that it is secreted during viral infection. Besides its subcellular localization, palmitoylation as a post-translational modification of the LC protein was predicted and further corroborated *in vitro*, with implications for its levels and subcellular localization. We also demonstrated the interaction of the LC protein with PDIA3, an important disulfide isomerase involved in the replication of other RNA viruses. We found that PDIs inhibition resulted in a change in subcellular localization and levels of the LC protein in FCV-infected cells. These results established some LC protein characteristics, indicating that it might play multiple roles in FCV-infected cells, consistent with its previously stated importance for a successful viral replication cycle.

## Results

### LC protein expression levels and subcellular localization during FCV infection

It has been previously established that the LC protein from FCV is translated late during infection as an LC-VP1 precursor protein processed by the viral protease-polymerase NS6/7 (Sosnovtsev et al., 1998). To determine the time post-infection at which the LC protein was first detected and to elucidate its expression levels during the FCV replicative cycle in CrFK cells, western blot assays were performed (Fig. 1). We found that the LC protein was first detected at 3 hours post-infection (hpi), and up to 9hpi, reaching its peak expression at 5 hpi and following a similar expression pattern as the VP1 protein (Fig. 1A and 1B).

**Figure 1.**
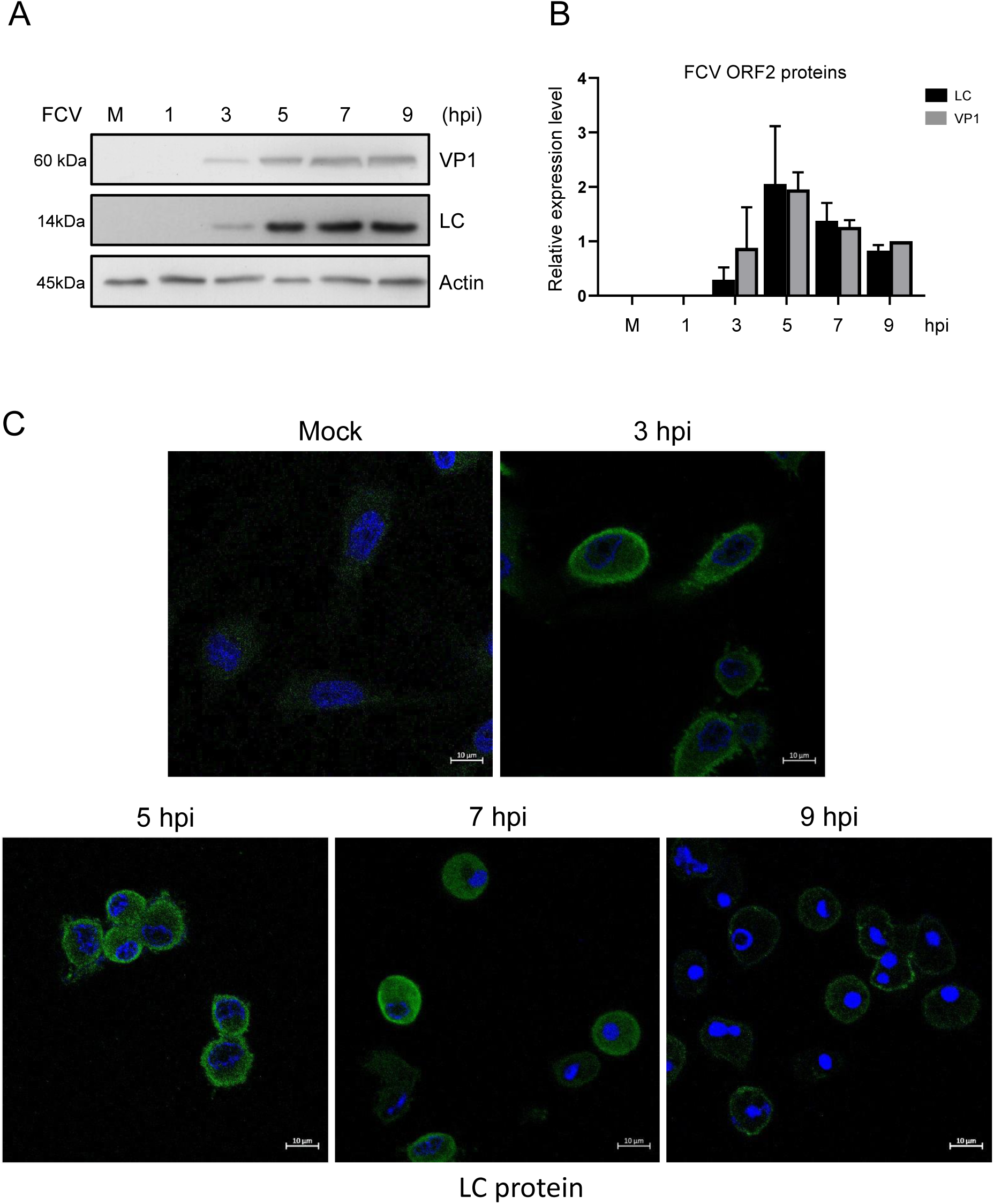
LC protein expression and subcellular localization through the FCV replicative cycle. A) Total protein extracts from mock-infected or FCV-infected cells at an MOI of 5 for 1, 3, 5, 7, and 9 h were obtained, and levels of LC and VP1 proteins were determined by western blotting. Actin was used as a loading control. B) LC and VP1 band intensities from scanned images were quantified using ImageJ software and shown as relative expression. Standard deviations were obtained from three independent experiments. Values of p≤0.0004 (***) calculated by T-test using GraphPad Prism 8.0 software are indicated. C) Mock-infected or FCV-infected cells at an MOI of 5 for 3, 5, 7, and 9 h were immunostained with an anti-LC serum (green). DAPI was used for nuclear (blue) staining. The cells were examined in a Zeiss LSM 700 confocal microscope. Images correspond to a z-stack of 15 slices and represent at least three independent experiments.

Once the expression levels of the LC protein were determined, we examined its subcellular distribution during infection. CrFK cells were infected with FCV at a multiplicity of infection (MOI) of 5, and the LC protein subcellular localization was analyzed at different times post-infection by confocal microscopy (Fig. 1C). At 3hpi, the LC protein was observed in both the cytoplasm and the periphery of the infected cells, which showed the first signs of cell rounding indicative of a cytopathic effect (Fig 1C). A similar subcellular localization was observed at 5 and 7hpi, but the signal intensity appeared greater than the observed at 3 hpi, correlating with the highest levels of the protein detected by western blotting (Fig. 1A and 1C). However, at 7 hpi, the LC protein was more evenly distributed throughout the cytoplasm, coinciding with established cytopathic effects and apoptosis (marked by pyknotic nuclei in infected cells) (Fig. 1C). At 9hpi, the LC protein signal was reduced, correlating with its detection by western blotting (Fig. 1A), and was mainly observed in the cell periphery in clusters that appeared to be located on the cellular membrane (Fig. 1C).

To further determine the specific subcellular localization of the LC protein in FCV-infected cells, we used subcellular markers of distinct compartments or organelles and analyzed colocalization with the LC protein by confocal microscopy at 5 hpi. To examine the possible localization of the LC protein in the mitochondria during infection, CrFK cells were infected at an MOI of 5 and its colocalization with the mitochondrial protein pyruvate dehydrogenase A1 (PDHA1) was determined by confocal microscopy (Fig. 2A). A discrete colocalization between the LC protein and PDHA1 was detected at 5 hpi (Fig. 2A). However, an evident localization of the exogenously expressed LC protein in CrFK cells transfected with the LC-pAm-Cyan, but not with the pAM-Cyan plasmid alone, as previously reported by our groupwork (Barrera-Vázquez et al., 2019), was confirmed by colocalization with the mitochondrial oxidoreductase coiled-coil-helix-coiled-coil-helix domain-containing protein 4 (CHCHD4), also known as MIA40, as observed in confocal microscopy assays (Supplementary Fig. 1). The discrete colocalization between PDHA1 and the LC protein in FCV-infected cells emphasizes the differences between the LC localization when expressed during infection and in a virus-free system, as has been reported for other viral proteins (Araujo et al., 2005; Boson et al., 2021; Pfaller et al., 2020).

**Figure 2.**
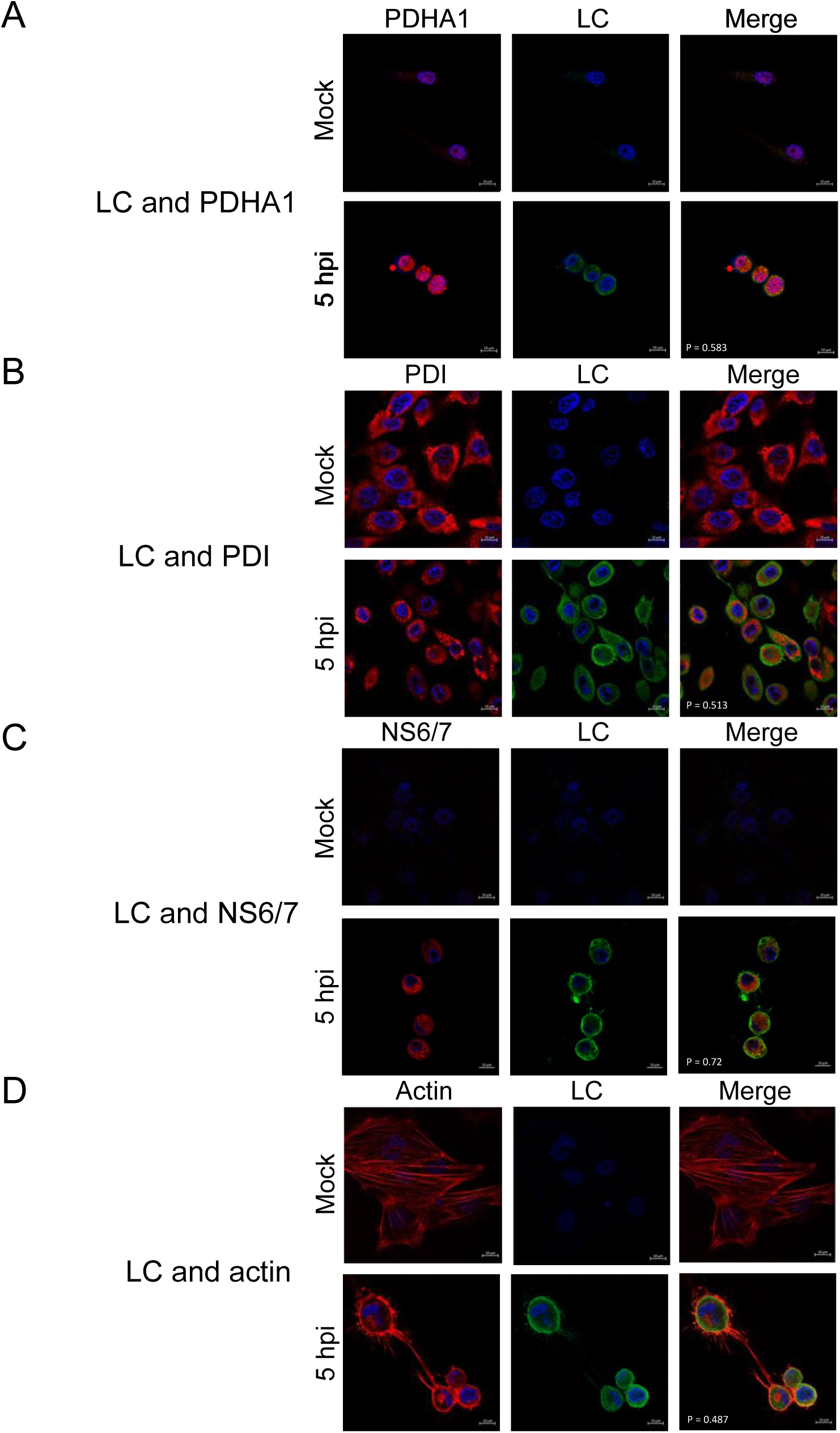
The LC protein is located in the cell periphery of FCV-infected cells. Mock-infected or FCV-infected cells at an MOI of 5 for 5 h were immunostained with an anti-LC serum (green) and A) an anti-PDHA1 (red), B) an anti-PDI (red), C) an anti-NS6/7 (red), and D) an anti-actin (red) antibodies. DAPI was used for nuclear (blue) staining. The cells were examined in a Zeiss LSM 700 confocal microscope. Images correspond to a z-stack of 15 slices and represent of at least three independent experiments. Merged images are indicated. Colocalization rates were calculated by Pearson’s coefficient correlation using Icy software (http://icy.biomageanalysis.org).

To determine if the LC protein was present in the endoplasmic reticulum (ER) during infection, its colocalization with the ER-resident lumen protein disulfide isomerase (PDI), used as a marker of FCV replication complexes (RC) (Cancio-Lonches et al., 2011), was analyzed by confocal microscopy (Fig. 2B). Again, a discrete colocalization was observed between these two proteins, suggesting that the LC protein is not present in this organelle. However, colocalization of the LC protein with the protease-polymerase NS6/7, another viral protein that is present in the RC as well as in the infected cytoplasm, was detected in discrete dots (Fig. 2C).

Since the LC protein is also observed in the periphery of the infected cells (Fig. 1B), it was possible that it could be associated with the cortical actin, an important component of the cell cytoskeleton and a key regulator of multiple cellular signaling pathways. Colocalization between the LC protein and actin was detected in the periphery of the infected cells. These results demonstrate that the LC protein expression increases steadily from 3 hpi, with a maximum expression at 5 hpi; is located near the RC, and at the cell periphery, most probably in association with cortical actin, but is not present in the mitochondria or the ER.

### LC of FCV is located at the inner and outer faces of the plasma membrane

Since the LC protein was found present in the cell periphery during different times of infection and seems to colocalize with the cortical actin, it is likely that this viral protein was also located on the outer face of the plasma membrane. To assess this possibility, its localization in non-permeabilized infected cells was determined by confocal microscopy at 5 hpi (Fig. 3A). The LC protein was found to be located at the outer face of the plasma membrane, but in focalized areas, in a capping-like pattern, contrasting its homogenous distribution on the entire inner face of the plasma membrane (Fig. 1C and 3A and 3B). These results demonstrate that besides its localization on the inner face of the cellular membrane, the LC protein is also focalized on the outer side of the plasma membrane.

**Figure 3.**
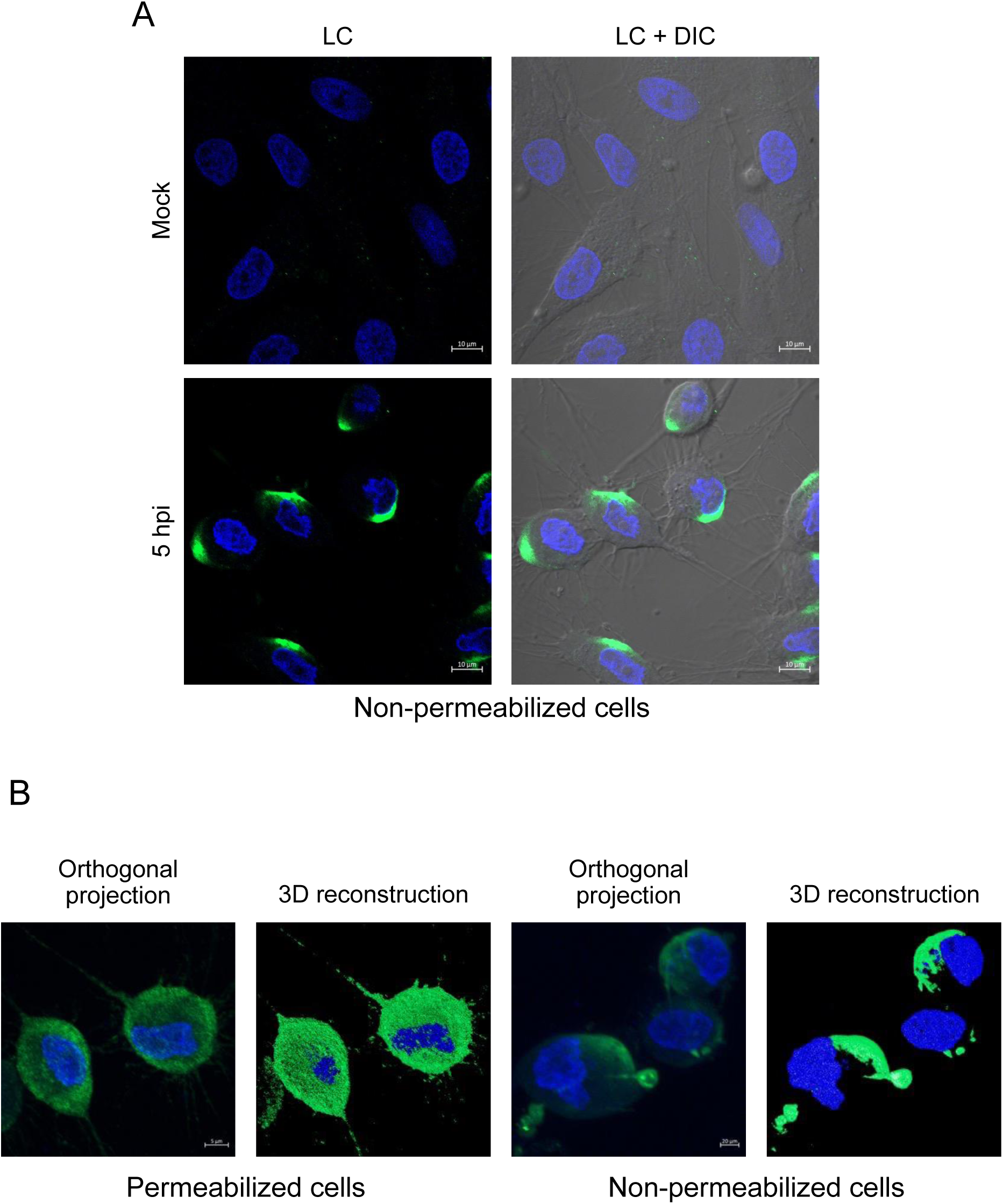
The LC protein is located on the outer face of the plasma membrane during FCV replication. A) Non-permeabilized, mock, or FCV-infected cells at an MOI of 5 for 5 h were immunostained with an anti-LC serum. The cells were examined in a Zeiss LSM 700 confocal microscope. Images correspond to a z-stack of 15 slices and reoresent at least three independent experiments. B) Orthogonal projections and 3D reconstructions from confocal microscopy assays of permeabilized and non-permeabilized FCV-infected cells.

### LC protein from FCV is palmitoylated in FCV-infected cells

Once the localization of LC protein was determined and its presence on the cell membrane was evidenced, we aimed to assess how this protein reaches this localization. A common determinant for proteins to locate to specific subcellular membranous compartments is through lipidation, which provides a hydrophobic anchor to a polypeptide embedded in its target membrane. Among lipidations, palmitoylation is a reversible post-translational modification carried on through the activity of the palmitoyl acyltransferases or zDHHC enzyme family members, directing specific subcellular localization of target proteins. There are multiple reports of viral proteins that are palmitoylated by the zDHHC palmitoyl transferases of their hosts, impacting their localization and/or stability (reviewed in Li et al., 2022; Veit, 2012).

To determine if the LC protein undergoes palmitoylation, we looked for putative palmitoylation signals in its primary structure using the palmitoylation prediction server GPS Palm V. 4.0 (Ning et al., 2021), and found that the LC protein contains four cysteine residues at positions 2, 5, 39, and 40 that could be S-palmitoylated. (Fig. 4A).

**Figure 4.**
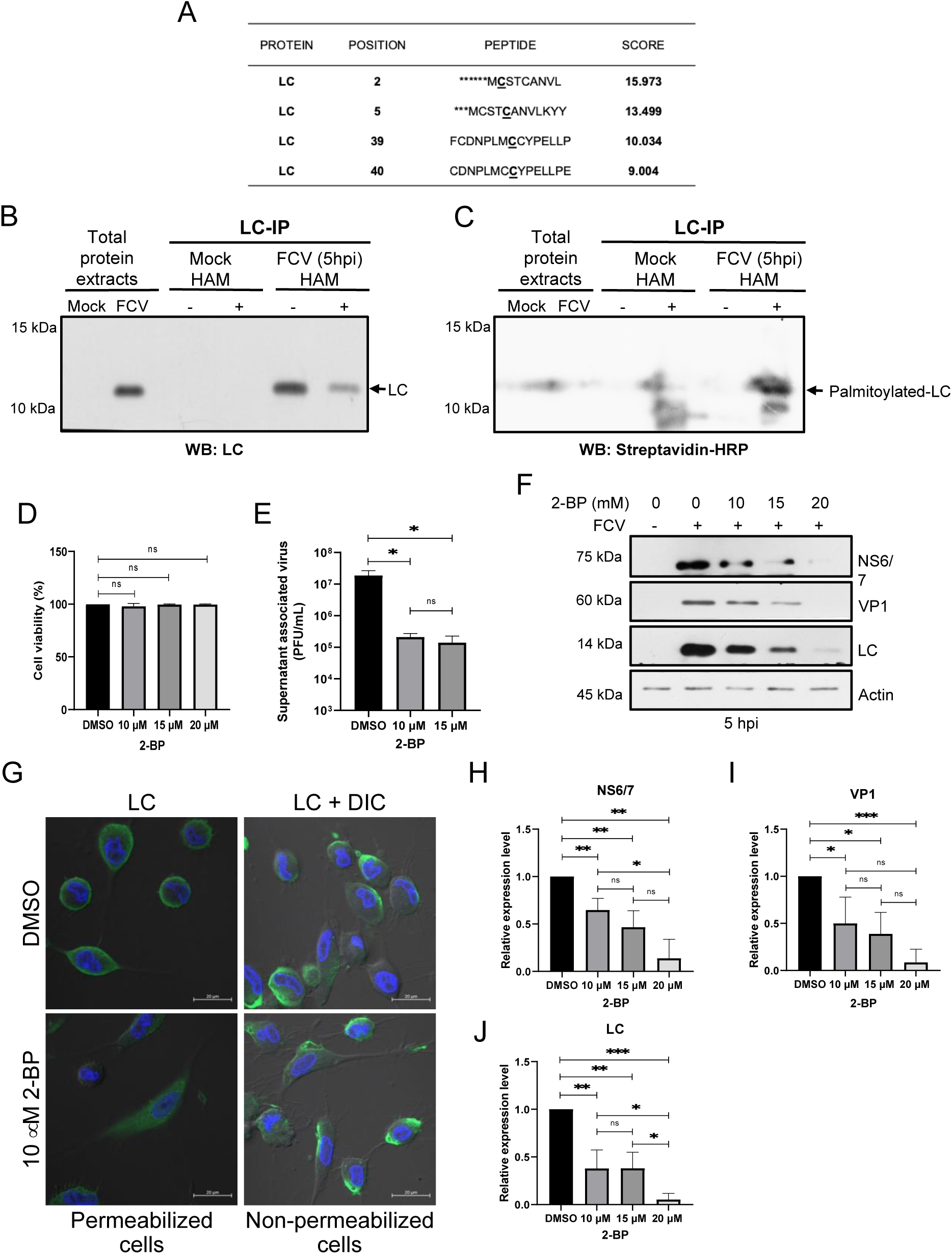
The LC protein form FCV is palmitoylated during infection. A) Putative cysteine residues susceptible to palmitoylation of the LC protein from FCV were identified using the GPS Palm 4.0 software. Total protein extracts from mock-infected or FCV-infected cells at an MOI of 5 for 5 h were obtained and subjected to immunoprecipitation with an anti-LC serum and subjected to ABE; the LC protein was detected by western blotting using B) anti-LC antibody or C) streptavidin-HRP. Hydroxylamine non-treated samples were used as an internal control. D) CrFK cells were treated with 10, 15, and 20 μM of 2-bromopalmitate and cell viability was assessed by an MTT assay. E) CrFK cells treated with 10 or 15 μM of 2-bromopalmitate were infected with FCV at an MOI of 5 for 5 h, and the supernatant-associated virus yield was quantified by plaque assay. Standard deviations were obtained from three independent experiments. Values of p≤ 0.05 (*) calculated by T-test using GraphPad Prism 8.0 software are indicated. F) Total protein extracts from cells treated with 10, 15, and 20 μM of 2-bromopalmitate, and mock-infected or infected with FCV at an MOI of 5 for 5 h were obtained, and levels of NS6/7, VP1, and LC proteins were determined by western blotting. Actin was used as a loading control. G) NS6/7, H) VP1, and I) LC band intensities from scanned images were quantified using ImageJ software and shown as relative expression. Standard deviations were obtained from three independent experiments. Values of p≤ 0.05 (*), p≤ 0.001 (**), and p≤ 0.0005 (***) calculated by T-test using GraphPad Prism 8.0 software are indicated. J) CrFK cells treated with DMSO or 10 μM 2-BP were infected with FCV at an MOI of 5 for 5 h, permeabilized or non-permeabilized, and immunostained with an anti-LC serum. The cells were examined in a Zeiss LSM 700 confocal microscope. Images correspond to a z-stack of 15 slices and represent at least three independent experiments.

To confirm that the LC protein was palmitoylated during FCV viral replication, it was immunoprecipitated from infected cell protein extracts and subjected to an Acyl-Biotin Exchange (ABE) assay (Fig. 4B). The specific immunoprecipitation of the LC protein from infected cells at 5 h was performed using an anti-LC serum, either in the presence or absence of hydroxylamine (HAM) and detected by western blotting using the anti-LC serum (Fig. 4B). A positive signal for the LC protein was found in the total protein extracts from infected cells and in the immunoprecipitated fraction from the extracts with or without HAM (Fig. 4B). To determine if the LC protein was palmitoylated, a parallel western blotting using streptavidin-HRP was performed (Fig. 4C). The immunoprecipitated LC protein from infected cell extracts treated with HAM, the only condition that allowed the ABE to occur, but not the immunoprecipitated LC protein from infected cell extracts not treated with HAM, was detected by streptavidin-HRP (Fig. 4C), indicating that this protein is palmitoylated during FCV replicative cycle.

To determine if the LC protein palmitoylation is required for efficient viral replication and could be related with its subcellular localization, CrFK cells were treated with different concentrations of the zDHHC palmitoyl transferases inhibitor 2-Bromopalmyate (2-BP) and infected with FCV at an MOI of 5 for 5 h. The supernatant viral titers, early and late viral protein production, and subcellular localization of LC protein were analyzed by plaque assays, western blotting, and immunofluorescence (Fig. 4D-4I). Viability of CrFK cells treated with 10, 15, and 20 mM of 2-BP for 5h was determined by MTT assays (Fig 4D), and no significant toxicity was observed at any of the concentrations used compared to the inhibitor vehicle (DMSO). Next, we analyzed the viral progeny present in the supernatant of drug-treated, FCV-infected CrFK cells. A 2-log reduction in viral titers from infected cells treated with 10 and 15 mM of 2-BP was found in comparison with the control vehicle-treated cells (Fig, 4E). To determine in which step of the viral cycle palmitoylation of the LC protein was required, the synthesis of early and late viral proteins in the presence or absence of 2-BP was analyzed by western blotting. A significant reduction of the early NS6/7 and late LC and VP1 viral proteins in cells infected for 5 h and treated with 10 and 15 mM of 2-BP was found compared to the infected DMSO-treated-cells. This suggests that the LC protein palmitoylation is required from the early stages for efficient FCV replication (Fig. 4F-4J).

Once we demonstrated that palmitoylation inhibition affected viral protein levels and virus yield, we wanted to determine if this post-translational modification plays a role in the localization of the LC protein to the cell periphery and the external face of the plasma membrane during infection. Therefore, infected CrFK cells treated with 10 mM 2-BP, a concentration at which a reduction but not a complete depletion of the LC protein levels was observed (Fig. 2F), showed a much more heterogeneous phenotype in which the typical cell-rounding phenotype was absent, and the LC protein was detected in the cytoplasm and to a lesser extent associated with the cortical actin (Fig. 4G). In contrast, no differences were observed between treated and non-treated, non-permeabilized cells regarding the LC protein localization outside of the cell membrane (Fig. 4G). These results suggest that the palmitoylation of the LC protein and cellular proteins influence, to some extent, the intracellular localization of the LC protein but is not required for its localization on the outer side of the plasma membrane. All these results taken together indicate that the LC protein is palmitoylated in FCV-infected cells and that palmitoylation influences the intracellular localization of the LC protein, but not its traffic to the outer face of the plasma membrane.

### LC protein is secreted to the extracellular medium

Given the localization of the LC protein at the cell periphery and the outer face of the plasma membrane in FCV-infected cells and its interaction with Annexin A2 (Abente et al., 2013), a protein involved in secretion, we investigated the possibility of the LC protein secretion during infection. The LC protein was detected in the supernatants of infected cells from 3hpi, the same time it was first detected by western blotting (Fig. 5A). This corresponds to the time at which the LC protein was first detected in a whole cell protein extract from infected cells (Fig. 1A). Moreover, the LC protein was also detected in increasing concentrations in the supernatants collected at 5, 7, and 9 hpi. To rule out the possibility that the presence of the LC protein in the supernatants was due to cell membrane disruption or cell lysis, a Sytox green entry assay was performed to evaluate cell permeation hourly up to 9 hpi using the microplate fluorometer Fluoroskan Ascent FL (Fig. 5B). No statistically significant changes in cell membrane permeation were observed at any of the analyzed times, including those in which the LC protein is located outside the infected cell and in the supernatant (Fig. 5B). Furthermore, no monomeric VP1 was detected in the supernatants of infected cells up to 7 hpi. These results strongly suggest that the LC protein is actively secreted during FCV replication cycle and may act as an extracellular viroporin.

**Figure 5.**
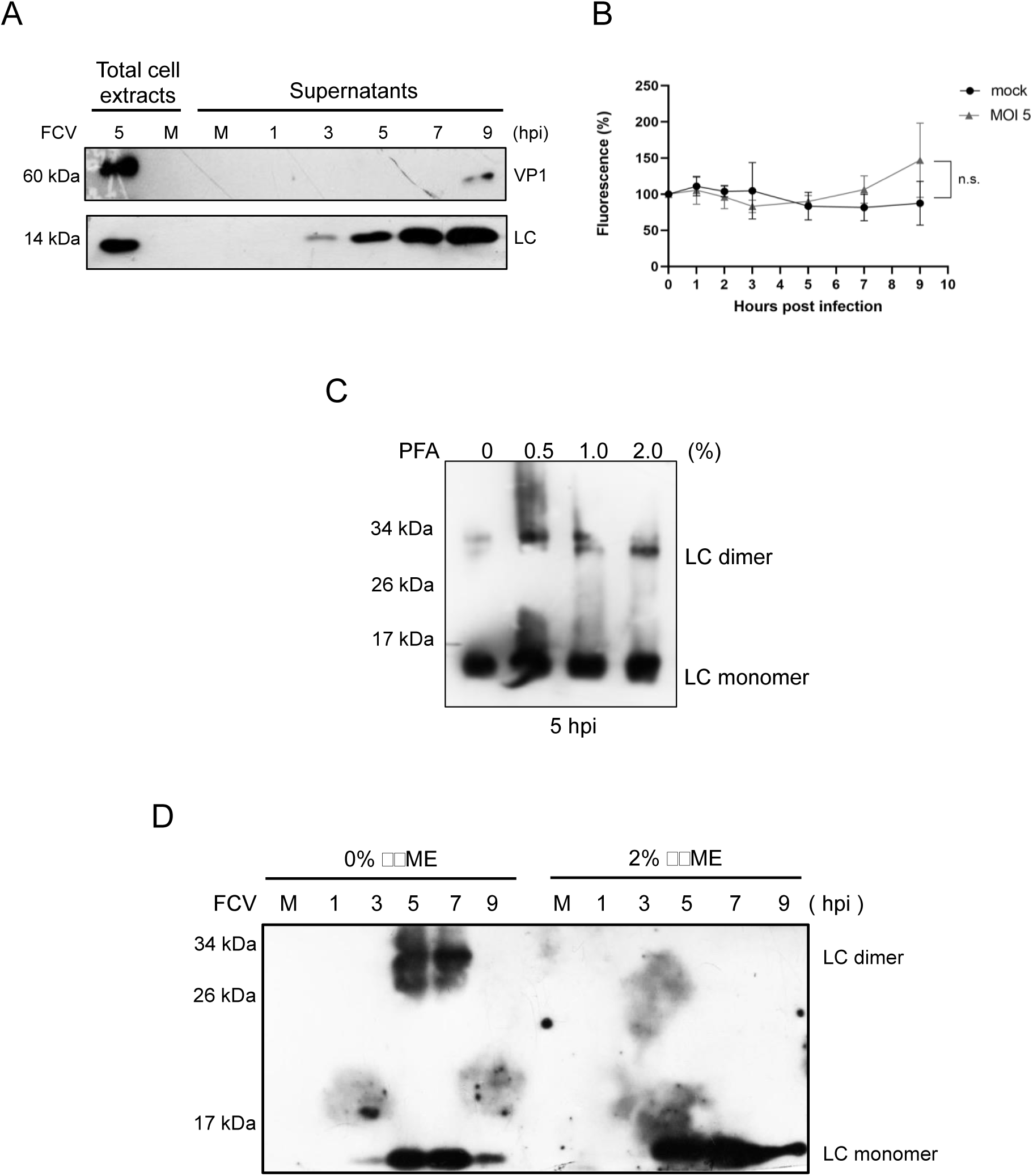
The LC protein can homodimerize and be secreted from FCV-infected cells. A) CrFK cells were mock-infected or FCV-infected at an MOI of 5 for 1, 3, 5, 7, and 9 h and the presence of the LC and VP1 proteins in the supernatants was determined by western blotting. Total protein extracts from mock-infected and FCV-infected cells at an MOI of 5 for 5 h were included as controls. B) Mock-infected and FCV-infected cells at an MOI of 5 for 5 h were incubated with Sytox Green for 10 min, and the total fluorescence was determined using the microplate fluorometer Fluoroskan Ascent FL. Standard deviations were obtained from three independent experiments. C) Total protein extracts form FCV-infected CrFK cells at an MOI of 5 for 5 h, untreated or treated with 0.5, 1, 10, and 2.0 % of FA were obtained, and the levels of the LC protein monomer and dimer were determined by western blotting. D) Total extracts from mock-infected and FCV-infected cells at 1, 3, 5, 7, and 9 h in the absence or presence of 2% β-ME were obtained and the levels of the LC protein monomer and dimer were determined by western blotting. LC protein monomers and dimers are indicated.

### LC from FCV forms homo-oligomers through disulfide bond formation

Viroporins often contain disulfide bonds that contribute to their structure, stability, or function (He et al., 2017; Holsinger & Alams, 1991; Hyser et al., 2010; Smertina et al., 2022). We have previously reported that purified Histag-LC protein homo-oligomerizes through disulfide bond formation; however, the oligomeric state of the LC protein in FCV-infected cells was still undetermined. To assess the oligomerization state of the LC protein during FCV replication cycle, infected cells at an MOI of 5 for 5 h were subjected to an *in vivo* cross-linking assay with increasing concentrations of paraformaldehyde (PFA) and the oligomeric state of the LC protein was analyzed by western blotting (Fig. 5C). A band of approximately 28 kDa, corresponding to a homo-dimeric form, was detected in the presence of PFA, suggesting the presence of LC homodimers in FCV infected cells (Fig. 5C). The dependance of LC protein oligomer formation on disulfide bonds was determined under reducing and non-reducing conditions using 2% β-mercaptoethanol (β−ME) as a reducing agent (Fig.5D). Again, in the non-reduced protein extracts, a band of 28 kDa corresponding to the homo-dimeric LC protein was detected at 5 and 7 hpi, while only the monomeric form was detected in β−ME-treated protein extracts collected at the same times post-infection (Fig. 5D). These results indicate that the LC protein forms homodimers through disulfide bonds.

### The LC protein from FCV interacts with the Protein Disulfide Isomerase A3

We have demonstrated that the LC protein forms disulfide bond-dependent homodimers in FCV-infected cells. Since we did not observe colocalization between the LC protein and the RE resident PDI and given the LC protein’s localization in the plasma membrane of infected cells, we hypothesized that PDIA3, abundantly present on both sides of the plasma membrane and involved in multiple roles in the replication cycle of diverse viruses (Mahmood et al., 2021), may interact with the LC protein. Colocalization between the LC protein and PDIA3 was observed mainly in the plasma membrane in permeabilized FCV-infected cells (Fig. 6A), demonstrating proximity between these two proteins at the cell periphery. Notably, higher levels of PDIA3 were observed on the plasma membrane of the FCV-infected, non-permeabilized cells, and in close contact with LC (PCC=0.622), than in mock-infected non-permeabilized cells (Fig. 6B). The association between the LC protein and PDI3A was corroborated by immunoprecipitation from cell lysates collected at 5 hpi and analyzed by western blotting under reducing and non-reducing conditions (Fig. 6C). PDI3A was detected in both mock-infected and FCV-infected total protein extracts, while the LC protein and VP1, included as a control, were detected only in infected cells (Fig. 6C). PDIA3 was co-immunoprecipitated with the LC protein, as it is strongly detected in the immunoprecipitation fraction from FCV-infected cells under reducing conditions, indicating its association with the LC protein during FCV-infection.

**Figure 6.**
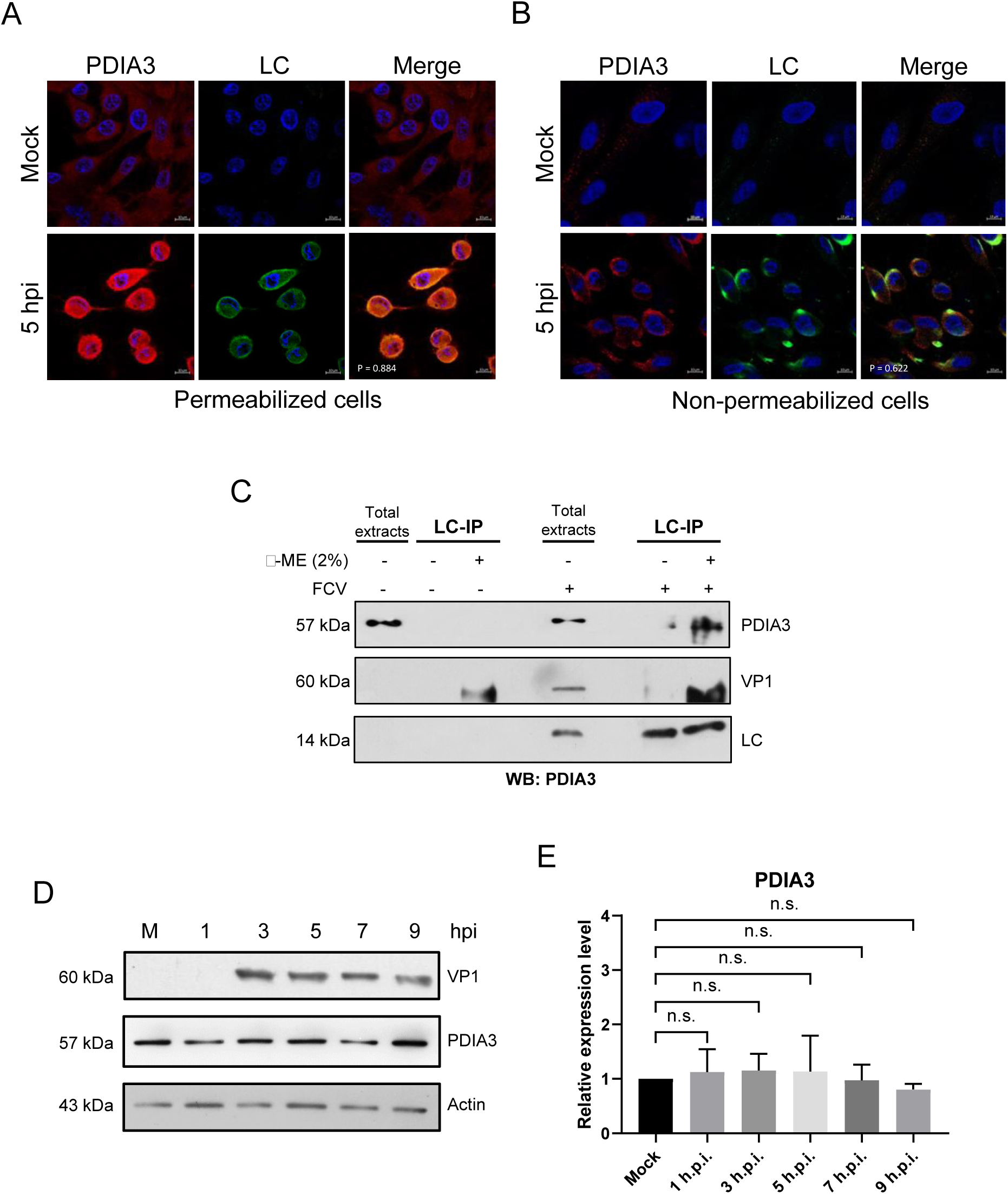
LC interacts with PDIA3 in FCV-infected cells. A) Permeabilized and B) non-permeabilized mock-infected and FCV-infected cells at an MOI of 5 for 5 h were immunostained with an anti-LC serum (green) and anti-PDIA3 (red) antibodies. DAPI was used for nuclear (blue) staining. The cells were examined in a Zeiss LSM 700 confocal microscope. Images correspond to a z-stack of 15 slices and represent at least three independent experiments. Merged images are indicated. Colocalization rates were calculated by Pearson’s coefficient correlation using Icy software (http://icy.biomageanalysis.org). C) Total protein extracts from mock-infected or FCV-infected cells at an MOI of 5 for 5 h were obtained and subjected to immunoprecipitation with an anti-LC serum. LC, VP1, and PDA3 proteins were detected by western blotting. Elutions were performed in the absence or presence of 2% ß-ME + 5 min heating at 95°C Total protein extracts from mock-infected and FCV-infected cells at 5 h were included as controls D) Total protein extracts from mock-infected or FCV-infected cells at an MOI of 5 for 1, 3, 5, 7, and 9 h were obtained, and levels of PDIA3 were determined by western blotting. VP1 indicates a virus infection. Actin was used as a loading control. E) PDIA3 band intensities from scanned images were quantified using ImageJ software and shown as relative expression. Standard deviations were obtained from three independent experiments.

To determine the relative expression of PDIA3 during cell infection, total protein extracts from mock-infected and FCV-infected cells at 1, 3, 5, 7, and 9 h were obtained, and the levels of PDIA3 were analyzed by western blotting (Fig. 6D). Similar PDIA3 levels were observed in both mock-infected and FCV-infected protein extracts at different times, indicating that this protein is not modulated during FCV replication. Taken together, these results demonstrate that the LC protein from FCV interacts with PDIA3, an important disulfide isomerase protein that participates in the replication cycle of various viruses and that PDIA3 levels are not modulated during FCV infection.

### PDIA3 protein is involved in the localization of the LC protein in the periphery of FCV-infected cells and in the formation of oligomers

To determine if PDIs have a role during FCV replicative cycle, their function was inhibited with the 16F16 compound (Hoffstrom et al., 2010). First, the viability of CrFK cells treated with 5, 10, 20, and 50µM concentrations of 16F16 for 5h was evaluated by MTT colorimetric assays (Fig. 7A). Since no changes in cell viability occurred in the presence of the 16F16 at the tested concentrations, the effect of PDIs inhibition in viral protein production was analyzed. Total protein extracts from 5, 10, 20, and 50 µM of 16F16-treated FCV-infected cells were obtained, and both VP1 and LC protein levels were analyzed by western blotting (Fig. 7B-D). A statistically significant reduction in the LC protein levels from 10 to 50 µM, was observed when compared to the DMSO control (Fig. 7C-D); moreover, a statistically significant reduction in the VP1 protein levels was observed at 20 and 50 µM of 16F16, suggesting that PDI functions are important for FCV replication.

**Figure 7.**
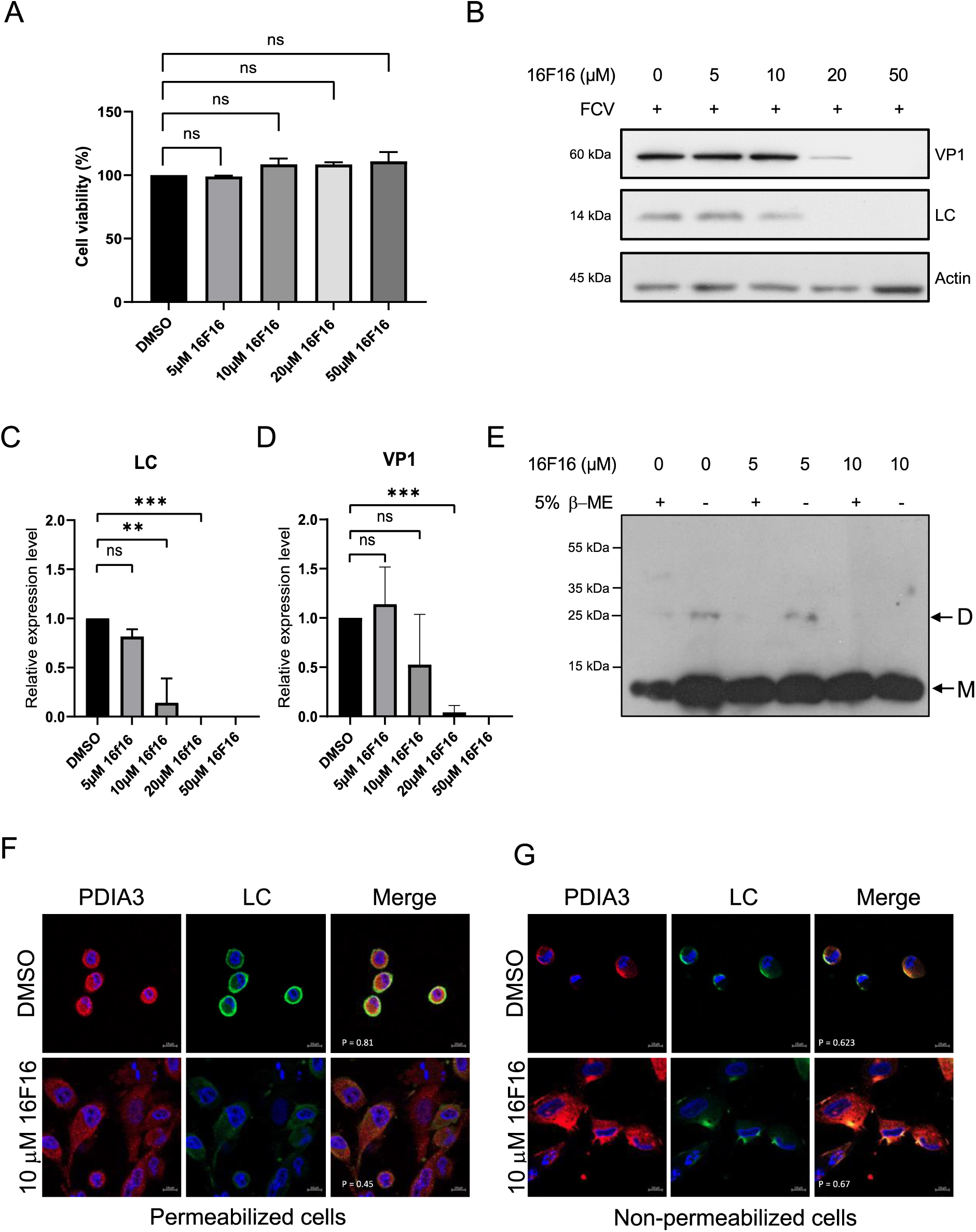
PDIs inhibitors reduce LC protein levels and oligomer formation and modify LC protein subcellular localization. A) CrFK cells were treated with 5, 10, 20, and 50 μM of 16F16, and cell viability was assessed by an MTT assay. B) Total protein extracts from mock-infected or FCV-infected cells at an MOI of 5 for 5 h, untreated or treated with 5, 10, 20, and 50 μM of 16F16 were obtained and levels of LC and VP1 proteins were determined by western blotting. Actin was used as a loading control. C) LC and D) VP1 band intensities from scanned images were quantified using ImageJ software and shown as relative expression. Standard deviations were obtained from three independent experiments. Values of p≤0.000 (***) calculated by T-test using GraphPad Prism 8.0 software are indicated. E) Total protein extracts from FCV-infected cells at an MOI of 5 for 5 h, untreated (0) or treated with 5 and 10 μM of 16F16 were obtained and analyzed by SDS-PAGE in the presence (+) or absence (-) of 5% β-ME. The presence of the LC protein monomers (M) and dimers (D) was determined by western blotting. F) Permeabilized and G) non-permeabilized mock-infected and FCV-infected cells at an MOI of 5 for 5 h and treated with DMSO or 10 μM of 16F16 were immunostained with an anti-LC serum (green) and anti-PDIA3 (red) antibodies. DAPI was used for nuclear (blue) staining. The cells were examined in a Zeiss LSM 700 confocal microscope. Images correspond to a z-stack of 15 slices and represent at least three independent experiments. Merged images are indicated. Colocalization rates were calculated by Pearson’s coefficient correlation using Icy software (http://icy.biomageanalysis.org).

To further analyze if PDIs function is involved in the LC protein subcellular localization and homodimer formation, 16F16-treated cells were infected with FCV and analyzed by immunofluorescence and western blotting (Fig. 7E-G). While DMSO-treated FCV-infected cells showed the typical cytopathic effect (Fig. 7F), no cell-rounding was observed in 10µM 16F16-treated infected cells (Fig. 7F). The LC protein signal is less intense in 16F16-treated cells in comparison to the DMSO-treated cells (Fig. 7F), in concordance with the reduction in LC protein levels observed in western blotting assays (Fig. 7B). Moreover, colocalization between the LC protein and PDIA3 (Pearson’s coefficient correlation of 0.81) was significantly decreased in the presence of 10 µM 16F16 (Pearson’s coefficient correlation of 0.45). To determine if the LC protein was able to reach the outer face of the plasma membrane when PDIs were inhibited, the LC protein localization was determined in non-permeabilized cells (Fig. 7G). Again, we found that the cytopathic effect in 16F16-treated infected cells was not as evident as in the DMSO-treated infected cells; however, the LC protein localization remained focalized in one side of the outer face of the plasma membrane and associated with PDIA3 in both 16F16 and DMSO-treated cells (Fig. 7F and 7FG), with Pearson’s correlation values of 0.623 and 0.67 respectively, suggesting that colocalization between PDIA3 and the LC protein is not affected by PDI inactivation. The role of PDIs in the LC protein oligomer formation was determined in infected cells treated with increasing concentrations of the PDI inhibitor 16F16. While the presence of the monomer (M) was observed under all conditions tested (Fig. 7E), dimer (D) formation was detected only at 0 and 5 μM 16F16. The lack of dimer formation at 10 μM 16F16 indicates that it requires the activity of a PDI. The 28 kDa band corresponding to the LC protein dimer was not observed in the presence of 5% β-ME, corroborating that they are formed through disulfide bonds. These results, taken together, suggest that PDIA3 plays a role in the LC protein stability and intracellular distribution during FCV replication and in the LC protein dimer formation but not in its translocation to the outer face of the plasma membrane from the infected cells.

## Discussion

The LC protein is unique to the members of the *Vesivirus* genus in the *Caliciviridae* family and is essential for successful viral replication. It is encoded in the ORF2 and expressed from the sgRNA as LC-VP1 precursor protein that is further processed by the viral protease-polymerase NS6/7, and it has not been found present in the mature viral particles. Although the LC protein from FCV was identified over 30 years ago, its importance in FCV infection was established by Abente *et al*. in 2013, who found that the whole protein is needed for a successful viral replication. They identified two conserved regions (CRs) necessary for its function and its interaction with the cellular factor Annexin A2 (Abente et al., 2013), which our workgroup further identified as being implicated in efficient FCV replication (Santos-Valencia et al., 2019). Even though it was identified that the LC protein participates in establishing the cytopathic effect, its function and mechanism of action still need to be fully elucidated.

The first clue suggesting that the LC protein from FCV acts as a viroporin came from our workgroup. Barrera *et al*. determined that the LC protein expression in a virus-free context resulted in its localization to the mitochondria, which induced periplasmic protein relocalization to the cytosol and triggered apoptosis, similar to other viroporins from RNA viruses (Madan et al., 2008). Furthermore, our workgroup reported that the purified LC recombinant protein forms homo-oligomers through disulfide bonds, and its expression is toxic in *E. coli* cells, likely through osmotic stress marked by plasmolytic bays formation, suggesting that the LC protein from FCV is a viroporin (Peñaflor-Téllez et al., 2020). We further hypothesized that disulfide bond formation in the LC protein may play a role in its toxicity and permeation capabilities, like the EboV Delta Peptide viroporin (He et al., 2017).

Viroporins, are virus-encoded transmembrane proteins that form pores or ionic channels and participate in multiple steps of viral replication in a virus-specific manner, making them attractive drug targets to minimize virus pathogenesis and replication (reviewed in Xia et al., 2022). Our previous works demonstrated that the LC protein from FCV has common viroporin characteristics (Peñaflor-Téllez 2022). In this work, we aimed to characterize the LC viroporin during FCV infection in cell culture. To this end, immunized mice serum was obtained to determine LC protein expression and subcellular localization throughout the FCV replication cycle. The LC protein expression from FCV was first detected by western blotting from 3 and up to 9hpi, with the highest levels achieved at 5 hpi, similar to the VP1 expression kinetics. When analyzing the subcellular localization of the LC protein in infected cells by immunofluorescence, we did not find it in the mitochondria, as when expressed in a virus-free system, which suggests that the LC protein may not play a direct role in the apoptosis induction through the intrinsic pathway, reported in FCV infection (Natoni et al., 2006; Roberts et al., 2003; Sosnovtsev et al., 2003). However, we cannot rule out an indirect role in apoptosis induction during infection. The main localization of the LC protein during infection was observed in the inner periphery of the infected cells, suggesting that this differential subcellular localization requires the presence of other viral proteins, has a post-translational modification or is the result of the infected cell context.

The localization of the LC protein from FCV on both the inner and outer face of the plasma membrane suggests that it could be involved in modulating cell surface signaling pathways. As was previously discussed by Abente et al. 2013, the CRII of the LC protein consists of conserved polyproline residues, which are common SH3 domain-binding motifs. Its colocalization with cytoskeletal protein actin at the inner face of the plasma membrane seen in some FCV-infected cells also suggests that the LC protein may be involved in cytoskeleton rearrangement, accordingly with its previously reported role in cytopathic effect establishment through its CRs (Abente et al., 2013). Whether the cytopathic effect is a consequence of the LC protein interaction with cytoskeletal proteins remains to be determined.

One-way proteins can be associated with the cell membrane is through lipidations as post-translational modifications. Although there were no previous reports of viral proteins in the *Caliciviridae* family undergoing this modification, our *in silico* prediction was the first hint suggesting that the LC viroporin could be palmitoylated during FCV infection. This was further demonstrated with the ABE method (Fig. 4), being LC the first caliciviral protein reported to be palmitoylated. The effect of the palmitoylation inhibition with 2-BP in FCV-infected cells resulted in a reduction of the LC protein levels. Furthermore, palmitoylation inhibition resulted in an alteration of the LC protein inner subcellular localization from the cell membrane to a more homogeneous cytoplasmic distribution. This result is similar to the changes that palmitoylated viral proteins undergo when this lipidation is inhibited, such as the TF protein of the Sindbis virus (SINV) (Ramsey et al., 2017) and the NS2 protein of the hepatitis C virus (HCV) (Wu et al., 2019), and even in other viroporins like ORF3 of hepatitis E virus (HEPEV) (Gouttenoire et al., 2018), and other viral proteins (reviewed by Li et al., 2022), suggesting that palmitoylation is involved in the LC protein localization to the inner face of the cell membrane. Interestingly, its localization to the external face of the membrane of FCV-infected cells in the absence or presence of 2-BP remained unchanged, even in the treated-infected cells that did not exhibit the typical cytopathic effect. This suggests that palmitoylation is primarily involved in the intracellular localization of the LC protein but not in the outer side of the cell membrane during FCV replication cycle. The palmitoyl-transferase enzyme(s) responsible for this lipidation, as well as the importance of this post-translational modification in the LC protein function inside the infected cell, remains to be determined.

As we found that the LC protein was focalized in the outer face of the cytoplasmic membrane of FCV-infected cells, we hypothesized that this protein was secreted during viral replication, similar to viroporins from other virus families like NSP4 from HuRoV (Tafazoli et al., 2001; Zhang et al., 2000) and Delta Peptide from EboV (He et al., 2017). The presence of the LC protein in the extracellular environment prior to apoptosis onset indicates that it is secreted and suggests that its function during FCV replication cycle is as an extracellular viroporin. This is the first report of a secreted calicivirus protein that has viroporin functional characteristics. Although the secretion of extracellular vesicles containing virions and viral RNA from FCV-infected cells has been previously reported (Mizenko et al., 2022), the role of free or vesicle-associated secreted FCV proteins has not been thoroughly studied, and it represents an attractive research line to understand the pathogenesis establishment of this virus better.

As the LC protein from FCV forms disulfide-dependent homo-oligomers in FCV-infected cells, we have hypothesized that it could occur through disulfide-isomerase proteins. PDI has been widely used as a marker of viral replication factories (Cancio-Lonches et al., 2011), but surprisingly, we found a discrete colocalization with the LC protein at 5hpi. However, the colocalization of the LC protein with PDIA3 at both sides of the cytoplasmic membrane and the confirmation of their interaction through co-immunoprecipitation strongly suggests that the LC protein from FCV interacts with this cellular chaperone for its correct folding/oligomerization. Inhibition of PDIs using 16F16 resulted in a reduction of the VP1 and LC protein levels and induced an alteration in the LC protein intracellular localization in FCV-infected cells, but not its focalized localization at the outer face of the plasma membrane. Moreover, the LC protein dimers’ formation depends on disulfide bonds through PDI’s function. The fact that the levels of both VP1 and LC proteins were reduced when PDIs function was inhibited by 16F16 with no effect on cell viability suggest that this compound (or other PDI inhibitors) could be used as an antiviral drug, as previously explored for other viruses (Fraternale et al., 2021). The downregulation of PDIA3 results in apoptosis triggering in several cell lines and is reported to be overexpressed in several cancer types, making it a potential biomarker (Tu et al., 2022) and drug target (Powell & Foster, 2021; Song et al., 2021). Therefore, PDI inhibition by 16F16 could also impact virus production through apoptosis modulation.

These results, taken together, elucidated the expression kinetics and subcellular localization of the LC protein as well as novel characteristics during the FCV replication cycle, such as its palmitoylation and association with PDIA3, that might be related to the LC protein dimer formation. Moreover, the fact that the LC protein is secreted during infection might be another viroporin characteristic, as has been reported for rotavirus NSP4 (Bugarcić & Taylor, 2006). The characterization of the LC protein in FCV-infected cells will provide new research lines to better understand this unique protein and its role during infection and will help to develop strategies to control and prevent FCV infection.

## Materials and methods

### Cell culture and viral infection

CrFK cells obtained from the American Type Culture Collection (ATCC) (Rockville, MD) were cultured in minimal essential medium (MEM) Advance supplemented with 5% fetal bovine serum (FBS) (Gibbco), 5000 U of penicillin, and 5 μg/ml of streptomycin. Cells were incubated at 37°C in 5% CO_2_. FCV Urbana strain was produced from the reverse genetic system using the pQ14 infectious clone kindly provided by Dr. Green (Laboratory of infectious diseases, NIAID, NIH, Bethesda, MD) (Sosnovtsev & Green, 1995).

Virus titers were quantified by plaque assay as described (Escobar-Herrera et al., 2007). The virus was diluted in MEM and used to infect 90% confluent CrFK cells at an MOI of 5, following two PBS washes (137 mM NaCl, 2.7 mM KCl, 10 mM Na_2_HPO_4_, and 1.8 mM KH_2_PO_4_) at 37°C for 5 min. Viral adsorption occurred over a 1-hour incubation at 37°C in 5% CO_2_, with gentle rocking every 15 min. After removing the viral inoculum, cells were washed with PBS at 37°C and incubated as previously described.

### Immunofluorescence assays

CrFK cells were grown on coverslips and infected as previously described. After infection and/or pharmacological treatment, cells were fixed with 3.7% paraformaldehyde (v/v) in PBS for 20 min and permeation by 0.1% Triton X-100 in PBS (v/v) for 5 min (permeabilization was omitted for non-permeabilized cells). Blocking was performed with 0.5% porcine skin gelatin for 30 min, followed by overnight (ON) incubation with the corresponding primary antibodies in PBS. Secondary Alexa 488 and CY5 fluorophores (Thermofisher Scientific) were diluted 1:200 in PBS and incubated 2 h at room temperature. Cells were stained with DAPI (4′,6-Diamidino-2-phenylindole dihydrochloride) (Thermofisher Scientific) in PBS for 5 min. Mounting was done with VectaShield (Vector laboratories), and analysis was performed using a Zeiss LSM 900 confocal microscope with ZEN lite software. All images were taken as optical sections in the z axis unless otherwise noted.

The antibodies for confocal microscopy and their respective dilutions were the following: anti-LC (mouse serum lab-generated) 1:120; anti-3CD (rabbit serum from Dr. Ian Goodfellow, Cambridge, UK) 1:80; anti-Annexin A2 (lab generated) 1:100; anti-PDIA3 (ABClonal Technology) 1:110; anti-PDI (ABClonal Technology) 1:100; anti-PDHA (ABClonal Technology) 1:150. Rhodamine-Phalloidin (Thermo Fisher Scientific) was used for actin staining according to the manufacturer’s instructions.

### Plasmid purification and transfection

CrFK cells grown on coverslips in 6-well plates (Corning) to 70% confluence were transfected with 3.5µg of pAm-Cyan and Wt-LC-pAm-Cyan plasmids (Barrera-Vázquez et al., 2019) using Lipofectamine 2000 (Thermofisher) in MEM medium (Gibco) following the standard transfection protocol. Transfected cells were then processed for immunofluorescence assays.

### Recombinant protein expression

Chemo-competent *E. coli* BL21(DE3)PlysS, transformed with pRSETA-LC plasmid (Peñaflor-Téllez 2022), were grown in LB medium (NaCl 0.17M, 1% peptone, 0.5% yeast extract) with ampicillin (100mg/µL) at 37°C and 200 rpm until O.D._600_ reached 0.4. Induction was done with IPTG (Isopropyl β-D-1-thiogalactopyranoside) (Thermofisher Scientific) at a final concentration of 0.1mM. Six hours post-induction, cells were sonicated following QIAexpressionist (Qiagen) protocols 9 and 10 using 8M UREA for protein denaturation. Membrane fractions were buffer-exchanged to PBS with Amicon Ultra 2-mL filters (Merck Millipore) and purified via FPLC with a Ni^2+^ column (Thermofisher).

### Animal immunization and serum purification

Bl6 mice were immunized with 50 µg of the His-tagged LC purified protein in PBS and TiterMax (Sigma-Aldrich) adjuvant (1:10 ratio) over four immunizations, each 14 days apart. Mice were euthanized via cardiac puncture, and serum was purified by differential centrifugation.

### Total protein extracts from CrFK cells

CrFK cells treated under specified conditions (infection, drug treatment or both), were harvested using plastic scrapers, pelleted at 1500 RCF for 5 min at 4°C, washed with pre-cooled PBS, and resuspended in RIPA buffer (150 mM NaCl, 1% Nonidet N-P40, 0.5% deoxycholate, 0.1% SDS, 50mM Tris-HCl pH 7.4). The lysates were incubated for 60 min at RT, centrifuged at 20,000 RCF for 30 min at 4°C, and the supernatants were collected. Protein degradation was minimized using cOmplete protease inhibitor cocktail (Roche), 5mM EDTA, and 0.1mM PMSF. Protein concentration was determined using a BCA protein kit assay (Thermo Scientific), and 10µg of protein was used per lane for western blotting. For non-reducing SDS-PAGE, Laemmli buffer lacked βME.

### SDS-PAGE and western blotting

SDS-PAGE followed the Bio-Rad Bulletin no. 6040 guide. Proteins were transferred to a nitrocellulose membrane (Bio-Rad) using Dunn buffer (10 mM NaCHO_3_, 3 mM Na_2_CO_3_, 8% methanol). Membranes were stained with Ponceau red, blocked with PBS-0.1% Tween 20 and 5% skimmed milk for 30 min at RT, and incubated with primary antibodies in PBS-Triton X100 (0.1%) (Sigma) ON at 4°C. Secondary antibodies conjugated with HRP (Jackson Immunoresearch) were diluted 1:10, 000 in PBS-Tween or PBS-Triton X100 with 1% skimmed milk for 2 h, at RT. Membranes were washed twice with PBS-Tween or Triton X-100 and developed using SuperSignal West Femto substrate (Thermo Scientific) and Carestream X-Ray films.

Primary antibodies used:

Α-Actin (Santa Cruz Biotechnology): 1:80,000. Anti-Vp1 (mouse serum, lab-generated): 1: 100,000. Anti-LC (mouse serum, lab-generated): 1:60,000. Anti-PDIA3 (ABClonal Technology): 1:10,000.

### *In situ* cell fixation for WB analysis

CrFK cells were infected and harvested by scrapping, centrifuged at 300 RCF for 5 min at 4°C, resuspended in ice-cold PBS, and centrifuged again. The supernatant was discarded, and cells were resuspended in PBS with PFA at the indicated concentrations, incubated with agitation for 20 min at RT, centrifuged, and resuspended in RIPA buffer for lysis and western blotting analysis.

### Pharmacological treatments

2-Bromopalmitate (2-BP) dissolved in DMSO at specified concentrations. Post-infection, cells were washed with PBS and incubated in MEM with 2% FBS and either 2-BP or 1% DMSO. For 16F16 (Santa Cruz Biotechnology), resuspended in DMSO to 0.311M and stored at -80°C, cells were treated similarly.

### Prediction of Palmitoylation Sites

S-Palmitoylation sites on the FCV LC protein were predicted using GPS-Palm 4.0 (https://gpspalm.biocuckoo.cn) in November 2022.

### Acyl-Biotin Exchange (ABE) assay

Detection of S-palmitoylation of the LC protein from FCV by ABE assay was performed as previously described (Brigidi & Bamji, 2013). Briefly, the cells were lysed in the presence of 50 mM N-ethylmaleimide (NEM) (Sigma) with lysis buffer containing 1% IGEPAL (Sigma), 50 mM Tris-HCl pH 7.5, 150 mM NaCl (Thermo Fisher), and 10% glycerol (Thermo Fisher). cOmplete protease inhibitor cocktail (Roche) and 0.1 mM phenylmethylsulfonyl fluoride (PMSF) (Sigma) were used as protease inhibitors. The target protein was immunoprecipitated with anti-LC serum for 2 h, at RT, and agarose beads coupled with protein A+G (Santa Cruz) were incubated at 4°C, ON under rocking conditions. The immunocomplex was incubated with 1 M hydroxylamine (HAM) (Sigma) in lysis buffer at pH 7.2 for 1 h at RT and washed 3 times. Biotinylation of the target protein was performed with Biotin-HPDP (Santa Cruz) in lysis buffer at pH 7.2 at 4°C for 1 h and washed 3 times. The samples were heated at 80°C for 10 min and centrifuged at 13,000 x g for 3 min to pellet the agarose beads. The samples were analyzed with sodium dodecyl sulfate-polyacrylamide gel electrophoresis (SDS-PAGE) and detected using Streptavidin-HRP (Thermo Fisher) or anti-LC serum as previously described.

### Membrane permeation assessment through Sytox Green entry assay

CrFK cells were seeded in 96-well culture plates until 80-90% confluence and mock-infected or infected with FCV at an MOI of 5. At the indicated times, 50 nM of Sytox Green diluted in a culture medium without SFB was added and incubated for 10 min before fluorescence measurement. The fluorescence intensity was determined at 1, 3, 5, 7, and 9 hpi with the fluorometer Fluoroskan Ascent FL (Thermo Fisher). Each measure was compared with the values obtained at 0 hpi.

### Protein precipitation from the cell supernatant

Protein precipitation was performed as previously described (Jakobs et al., 2013). Briefly, supernatants of CrFK cells, mock-infected or infected with FCV at a MOI of 5, were collected, and cell debris was removed by centrifugation at 500 RCF at 4°C for 5 min, discarding the pellet, and at 2,000 RCF at 4°C for 30 min, discarding the pellet. Finally, centrifugation at 20,000 RCF at 4°C for 30 min was performed, keeping the supernatants. Five hundred µL of methanol (J.T. Baker) and 125 µL of chloroform (J. T. Baker) were added to 500 µL of supernatants and mixed by vortexing until a homogeneous solution was formed. The samples were centrifuged at 13,000 RCF for 5 min. The aqueous phase was carefully removed using a vacuum pump, and 500 µL of methanol was added before mixing by vortexing until the protein pellet was broken down. The samples were centrifuged at 13,000 RCF for 5 min, the aqueous phase was discarded using a vacuum pump, and the pellet was resuspended in RIPA lysis buffer (10mM Tris-Cl, 1mM EDTA, 0.5mM EGTA, 1% Triton x100, 0.1% sodium deoxycholate, 0.1% SDS, 140 mM NaCl at pH 8) with cOmplete protease inhibitor cocktail (Roche) for SDS-PAGE and western blotting analysis.

### LC co-immunoprecipitation assays

CrFK cells were seeded, grown, and infected as previously described. Seven x 10^7^ cells were harvested with a scraper and washed once with ice-cold PBS before RIPA lysis and protein extraction, as previously described. Total protein extracts were incubated with agitation with 2.3 µl α-LC mouse serum for 2 h at RT. Nine µl of protein A/G plus agarose beads (Santa Cruz Biotechnology) was added and incubated at 4°C ON. Samples were centrifuged at 500 RCF at 4°C for 5 min and washed 2 times with PBS. Protein was eluted in Laemmli buffer 2X at 95°C for 5 min, for SDS-PAGE and western blotting analysis.

### MMT Assay

CrFK cells were plated in 96-well culture plates (10,000 cells/well) in Dulbecco’s modified Eagle’s medium (DMEM) supplemented with 5% FBS until 90% confluence. Drug treatment was given according to the experimental requirement. The medium was removed, and 100 μL of fresh medium was added with 10 MTT (Sigma) to a final concentration of 1.2 mM; the cells were incubated for 4 h at 37° C and protected from light. The formazan crystals formed by the cells were dissolved by adding 100 µL of 5% SDS (Bio-Rad) in 1N HCl (Thermo Fisher) and incubated for 12 h at 37°C. Finally, the absorbance was measured at 540 nm using the microplate reader.

## Acknowledgements.

This work was supported by a grant from Consejo Nacional de Humanidades, Ciencias y Tecnologías (Conahcyt), [Proyecto 302965, PRONAII 3: Infecciones virales del tracto gastrointestinal]. YPT, JGM, CEMR, EICM, and CPI received a scholarship from Conahcyt. The funders had no role in study design, data collection, and interpretation, or the decision to submit the work for publication. We thank Juan Ludert for their critical comments on the manuscript.

## Contributor roles

Conceptualization: YPT, and ALGE.

Formal analysis: YPT, JGM, EICM, CPI, and ALGE.

Funding acquisition: ALGE.

Investigation and Methodology: YPT, JGM, EICM, CEMR, CPI, and ALGE.

Supervision: ALGE.

Validation: YPT, JGM, EICM, CPI, and ALGE.

Writing original draft: YPT.

Writing, review, and editing: YPT and ALGE.

## Supplementary

**Supplementary figure 1.**
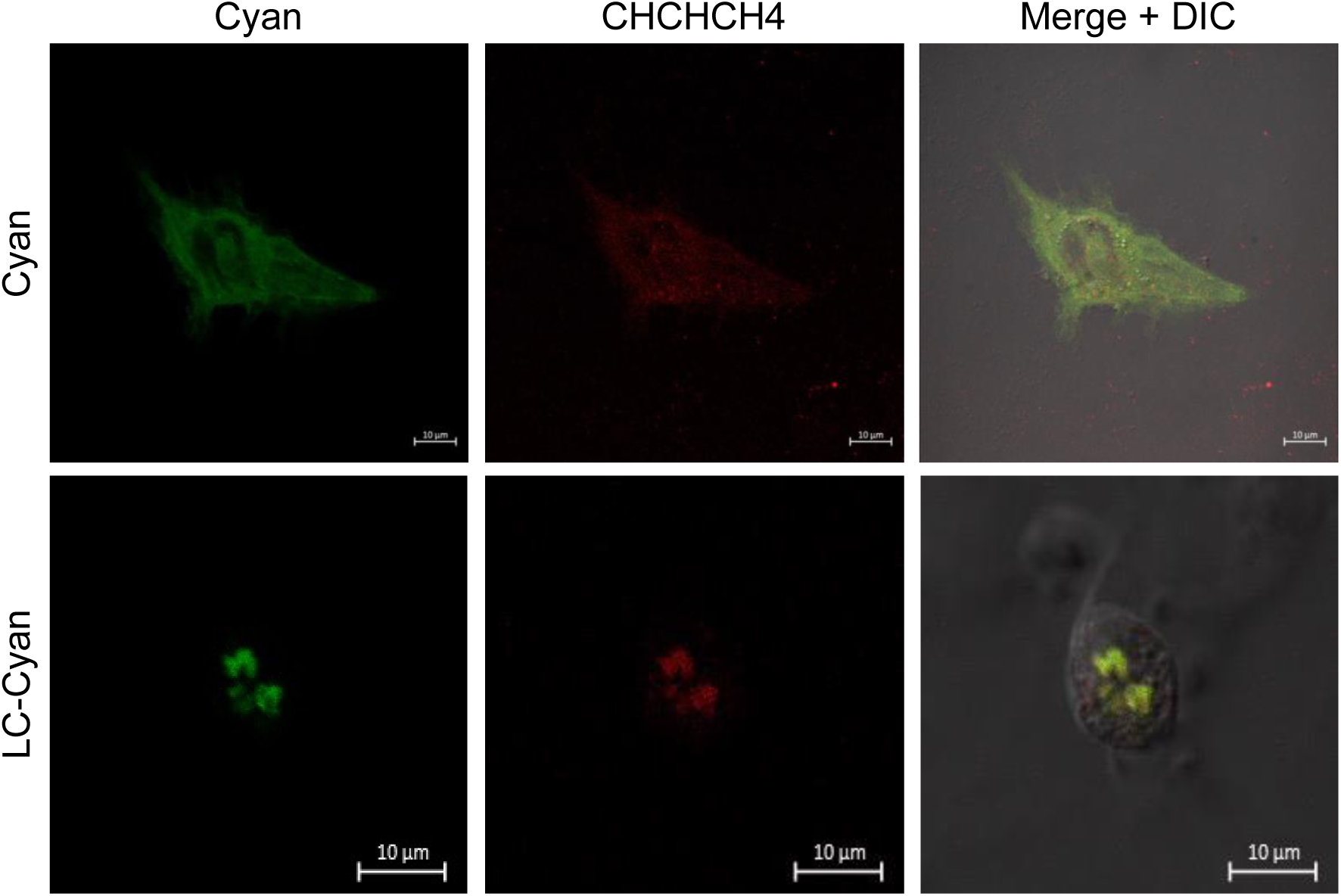
Mitochondrial localization of the LC-Cyan protein in transfected cells. CrFK cells were transfected with Cyan or Cyan-LC proteins for 48 h and immunostained with an anti-LC serum (green) and anti-CHCHCH4 (red) antibodies. The cells were examined in a Zeiss LSM 700 confocal microscope. Images correspond to a z-stack of 15 slices and represent at least three independent experiments. Merged images are indicated.

## Bibliography

1. Abente, E. J., Sosnovtsev, S. V., Sandoval-Jaime, C., Parra, G. I., Bok, K., & Green, K. Y. (2013). The Feline Calicivirus Leader of the Capsid Protein Is Associated with Cytopathic Effect. Journal of Virology, 87(6), 3003–3017. 10.1128/jvi.02480-12

2. Araujo, F. D., Stracker, T. H., Carson, C. T., Lee, D. V., & Weitzman, M. D. (2005). Adenovirus Type 5 E4orf3 Protein Targets the Mre11 Complex to Cytoplasmic Aggresomes. Journal of Virology, 79(17), 11382–11391. 10.1128/JVI.79.17.11382-11391.2005

3. Barrera-Vázquez, O. S., Cancio-Lonches, C., Hernández-González, O., Chávez-Munguia, B., Villegas-Sepúlveda, N., & Gutiérrez-Escolano, A. L. (2019). The feline calicivirus leader of the capsid protein causes survivin and XIAP downregulation and apoptosis. Virology, 527(November 2018), 146–158. 10.1016/j.virol.2018.11.017

4. Bergmann, M., Speck, S., Rieger, A., Truyen, U., & Hartmann, K. (2019). Antibody response to feline calicivirus vaccination in healthy adult cats. Viruses, 11(8), 1–14. 10.3390/v11080702

5. Bordicchia, M., Fumian, T. M., Van Brussel, K., Russo, A. G., Carrai, M., Le, S.-J., Pesavento, P. A., Holmes, E. C., Martella, V., White, P., Beatty, J. A., Shi, M., & Barrs, V. R. (2021). Feline Calicivirus Virulent Systemic Disease: Clinical Epidemiology, Analysis of Viral Isolates and In Vitro Efficacy of Novel Antivirals in Australian Outbreaks. Viruses, 13(10), 2040. 10.3390/v13102040

6. Boson, B., Legros, V., Zhou, B., Siret, E., Mathieu, C., Cosset, F.-L., Lavillette, D., & Denolly, S. (2021). The SARS-CoV-2 envelope and membrane proteins modulate maturation and retention of the spike protein, allowing assembly of virus-like particles. Journal of Biological Chemistry, 296, 100111. 10.1074/jbc.RA120.016175

7. Brigidi, G. S., & Bamji, S. X. (2013). Detection of Protein Palmitoylation in Cultured Hippocampal Neurons by Immunoprecipitation and Acyl-Biotin Exchange (ABE). Journal of Visualized Experiments, 72. 10.3791/50031

8. Bugarcić, A., & Taylor, J. A. (2006). Rotavirus Nonstructural Glycoprotein NSP4 Is Secreted from the Apical Surfaces of Polarized Epithelial Cells. Journal of Virology, 80(24), 12343–12349. 10.1128/jvi.01378-06

9. Cancio-Lonches, C., Yocupicio-Monroy, M., Sandoval-Jaime, C., Galvan-Mendoza, I., Ureña, L., Vashist, S., Goodfellow, I., Salas-Benito, J., & Gutiérrez-Escolano, A. L. (2011). Nucleolin Interacts with the Feline Calicivirus 3′ Untranslated Region and the Protease-Polymerase NS6 and NS7 Proteins, Playing a Role in Virus Replication. Journal of Virology, 85(16), 8056–8068. 10.1128/JVI.01878-10

10. Carter, M. J., Milton, I. D., Turner, P. C., Meanger, J., Bennett, M., & Gaskell, R. M. (1992). Identification and sequence determination of the capsid protein gene of feline calicivirus. Archives of Virology, 122(3–4), 223–235. 10.1007/BF01317185

11. Deschamps, J.-Y., Topie, E., & Roux, F. (2015). Nosocomial feline calicivirus-associated virulent systemic disease in a veterinary emergency and critical care unit in France. Journal of Feline Medicine and Surgery Open Reports, 1(2), 205511691562158. 10.1177/2055116915621581

12. Escobar-Herrera, J., Medina-Ramírez, F. J., & Gutiérrez-Escolano, A. L. (2007). A carboxymethyl-cellulose plaque assay for feline calicivirus. Journal of Virological Methods, 146(1–2), 393–396. 10.1016/j.jviromet.2007.07.013

13. Fraternale, A., Zara, C., De Angelis, M., Nencioni, L., Palamara, A. T., Retini, M., Di Mambro, T., Magnani, M., & Crinelli, R. (2021). Intracellular Redox-Modulated Pathways as Targets for Effective Approaches in the Treatment of Viral Infection. International Journal of Molecular Sciences, 22(7), 3603. 10.3390/ijms22073603

14. Gouttenoire, J., Pollán, A., Abrami, L., Oechslin, N., Mauron, J., Matter, M., Oppliger, J., Szkolnicka, D., Dao Thi, V. L., van der Goot, F. G., & Moradpour, D. (2018). Palmitoylation mediates membrane association of hepatitis E virus ORF3 protein and is required for infectious particle secretion. PLoS Pathogens, 14(12), 1–24. 10.1371/journal.ppat.1007471

15. He, J., Melnik, L. I., Komin, A., Wiedman, G., Fuselier, T., Morris, C. F., Starr, C. G., Searson, P. C., Gallaher, W. R., Hristova, K., Garry, R. F., & Wimley, W. C. (2017). Ebola Virus Delta Peptide Is a Viroporin. Journal of Virology, 91(16), 1–14. 10.1128/jvi.00438-17

16. Herbert, T. P., Brierley, I., & Brown, T. D. K. (1996). Detection of the ORF3 polypeptide of feline calicivirus in infected cells and evidence for its expression from a single, functionally bicistronic, subgenomic mRNA. Journal of General Virology, 77(1), 123–127. 10.1099/0022-1317-77-1-123

17. Hoffstrom, B. G., Kaplan, A., Letso, R., Schmid, R. S., Turmel, G. J., Lo, D. C., & Stockwell, B. R. (2010). Inhibitors of protein disulfide isomerase suppress apoptosis induced by misfolded proteins. Nature Chemical Biology, 6(12), 900–906. 10.1038/nchembio.467

18. Hofmann-Lehmann, R., Hosie, M. J., Hartmann, K., Egberink, H., Truyen, U., Tasker, S., Belák, S., Boucraut-Baralon, C., Frymus, T., Lloret, A., Marsilio, F., Pennisi, M. G., Addie, D. D., Lutz, H., Thiry, E., Radford, A. D., & Möstl, K. (2022). Calicivirus Infection in Cats. Viruses, 14(5), 937. 10.3390/v14050937

19. Holsinger, L. J., & Alams, R. (1991). Influenza virus M2 integral membrane protein is a homotetramer stabilized by formation of disulfide bonds. Virology, 183(1), 32–43. 10.1016/0042-6822(91)90115-R

20. Huang, C., Hess, J., Gill, M., & Hustead, D. (2010). A dual-strain feline calicivirus vaccine stimulates broader cross-neutralization antibodies than a single-strain vaccine and lessens clinical signs in vaccinated cats when challenged with a homologous feline calicivirus strain associated with virulent sys. Journal of Feline Medicine and Surgery, 12(2), 129–137. 10.1016/j.jfms.2009.08.006

21. Hurley, K. F., Pesavento, P. A., Pedersen, N. C., Poland, A. M., Wilson, E., & Foley, J. E. (2004). An outbreak of virulent systemic feline calicivirus disease. Journal of the American Veterinary Medical Association, 224(2), 241–249. 10.2460/javma.2004.224.241

22. Hyser, J. M., Collinson-Pautz, M. R., Utama, B., & Estes, M. K. (2010). Rotavirus disrupts calcium homeostasis by NSP4 viroporin activity. MBio, 1(5), 1–12. 10.1128/mBio.00265-10

23. Jakobs, C., Bartok, E., Kubarenko, A., Bauernfeind, F., & Hornung, V. (2013). Immunoblotting for Active Caspase-1 (pp. 103–115). 10.1007/978-1-62703-523-1_9

24. Lesbros, C., Martin, V., Najbar, W., Sanquer, A., McGahie, D., Eun, H. M., & Gueguen, S. (2013). Protective efficacy of the calicivirus valency of the Leucofeligen vaccine against a virulent heterologous challenge in kittens. Veterinary Medicine International, 2013. 10.1155/2013/232397

25. Li, X., Shen, L., Xu, Z., Liu, W., Li, A., & Xu, J. (2022). Protein Palmitoylation Modification During Viral Infection and Detection Methods of Palmitoylated Proteins. Frontiers in Cellular and Infection Microbiology, 12. 10.3389/fcimb.2022.821596

26. Madan, V., Castelló, A., & Carrasco, L. (2008). Viroporins from RNA viruses induce caspase-dependent apoptosis. Cellular Microbiology, 10(2), 437–451. 10.1111/j.1462-5822.2007.01057.x

27. Mahmood, F., Xu, R., Awan, M. U. N., Song, Y., Han, Q., Xia, X., & Zhang, J. (2021). PDIA3: Structure, functions and its potential role in viral infections. Biomedicine & Pharmacotherapy, 143, 112110. 10.1016/j.biopha.2021.112110

28. Mizenko, R. R., Brostoff, T., Jackson, K., Pesavento, P. A., & Carney, R. P. (2022). Extracellular Vesicles (EVs) Are Copurified with Feline Calicivirus, yet EV-Enriched Fractions Remain Infectious. Microbiology Spectrum, 10(4). 10.1128/spectrum.01211-22

29. Pedersen, N. C., Elliott, J. B., Glasgow, A., Poland, A., & Keel, K. (2000). An isolated epizootic of hemorrhagic-like fever in cats caused by a novel and highly virulent strain of feline calicivirus. Veterinary Microbiology, 73(4), 281–300. 10.1016/S0378-1135(00)00183-8

30. Peñaflor-Téllez, Y., Chávez-Munguía, B., Lagunes-Guillén, A., Salazar-Villatoro, L., & Gutiérrez-Escolano, A. L. (2022). The Feline Calicivirus Leader of the Capsid Protein Has the Functional Characteristics of a Viroporin. Viruses, 14(3), 635. 10.3390/v14030635

31. Peñaflor-Téllez, Y., Miguel-Rodríguez, C. E., & Gutiérrez-Escolano, A. L. (2020). The Caliciviridae Family. In Reference Module in Biomedical Sciences. Elsevier. 10.1016/b978-0-12-818731-9.00027-6

32. Pfaller, C. K., Bloyet, L.-M., Donohue, R. C., Huff, A. L., Bartemes, W. P., Yousaf, I., Urzua, E., Clavière, M., Zachary, M., de Masson d’Autume, V., Carson, S., Schieferecke, A. J., Meyer, A. J., Gerlier, D., & Cattaneo, R. (2020). The C Protein Is Recruited to Measles Virus Ribonucleocapsids by the Phosphoprotein. Journal of Virology, 94(4). 10.1128/JVI.01733-19

33. Powell, L. E., & Foster, P. A. (2021). Protein disulphide isomerase inhibition as a potential cancer therapeutic strategy. Cancer Medicine, 10(8), 2812–2825. 10.1002/cam4.3836

34. Ramsey, J., Renzi, E. C., Arnold, R. J., Trinidad, J. C., & Mukhopadhyay, S. (2017). Palmitoylation of Sindbis Virus TF Protein Regulates Its Plasma Membrane Localization and Subsequent Incorporation into Virions. Journal of Virology, 91(3). 10.1128/JVI.02000-16

35. Santos-Valencia, J. C., Cancio-Lonches, C., Trujillo-Uscanga, A., Alvarado-Hernández, B., Lagunes-Guillén, A., & Gutiérrez-Escolano, A. L. (2019). Annexin A2 associates to feline calicivirus RNA in the replication complexes from infected cells and participates in an efficient viral replication. Virus Research, 261(June 2018), 1–8. 10.1016/j.virusres.2018.12.003

36. Smertina, E., Carroll, A. J., Boileau, J., Emmott, E., Jenckel, M., Vohra, H., Rolland, V., Hands, P., Hayashi, J., Neave, M. J., Liu, J.-W., Hall, R. N., Strive, T., & Frese, M. (2022). Lagovirus Non-structural Protein p23: A Putative Viroporin That Interacts With Heat Shock Proteins and Uses a Disulfide Bond for Dimerization. Frontiers in Microbiology, 13. 10.3389/fmicb.2022.923256

37. Song, D., Guo, M., Wu, K., Hao, J., Nie, Y., & Fan, D. (2021). Silencing of ER-resident oxidoreductase PDIA3 inhibits malignant biological behaviors of multidrug-resistant gastric cancer. Acta Biochimica et Biophysica Sinica, 53(9), 1216–1226. 10.1093/abbs/gmab101

38. Sosnovtsev, S., & Green, K. Y. (1995). RNA Transcripts Derived from a Cloned Full-Length Copy of the Feline Calicivirus Genome Do Not Require VpG for Infectivity. Virology, 210(2), 383–390. 10.1006/viro.1995.1354

39. Sosnovtsev, S. V., & Green, K. Y. (2000). Identification and genomic mapping of the ORF3 and VPg proteins in feline calicivirus virions. Virology, 277(1), 193–203. 10.1006/viro.2000.0579

40. Sosnovtsev, S. V., Sosnovtseva, S. A., & Green, K. Y. (1998a). Cleavage of the Feline Calicivirus Capsid Precursor Is Mediated by a Virus-Encoded Proteinase. Journal of Virology, 72(4), 3051–3059. 10.1128/JVI.72.4.3051-3059.1998

41. Sosnovtsev, S. V., Sosnovtseva, S. A., & Green, K. Y. (1998b). Cleavage of the Feline Calicivirus Capsid Precursor Is Mediated by a Virus-Encoded Proteinase. Journal of Virology, 72(4), 3051–3059. 10.1128/jvi.72.4.3051-3059.1998

42. Tafazoli, F., Zeng, C. Q., Estes, M. K., Magnusson, K.-E., & Svensson, L. (2001). NSP4 Enterotoxin of Rotavirus Induces Paracellular Leakage in Polarized Epithelial Cells. Journal of Virology, 75(3), 1540–1546. 10.1128/jvi.75.3.1540-1546.2001

43. Tu, Z., Ouyang, Q., Long, X., Wu, L., Li, J., Zhu, X., & Huang, K. (2022). Protein Disulfide-Isomerase A3 Is a Robust Prognostic Biomarker for Cancers and Predicts the Immunotherapy Response Effectively. Frontiers in Immunology, 13. 10.3389/fimmu.2022.837512

44. Veit, M. (2012). Palmitoylation of virus proteins. Biology of the Cell, 104(9), 493–515. 10.1111/boc.201200006

45. Wu, M.-J., Shanmugam, S., Welsch, C., & Yi, M. (2019). Palmitoylation of Hepatitis C Virus NS2 Regulates Its Subcellular Localization and NS2-NS3 Autocleavage. Journal of Virology, 94(1). 10.1128/JVI.00906-19

46. Xia, X., Cheng, A., Wang, M., Ou, X., Sun, D., Mao, S., Huang, J., Yang, Q., Wu, Y., Chen, S., Zhang, S., Zhu, D., Jia, R., Liu, M., Zhao, X.-X., Gao, Q., & Tian, B. (2022). Functions of Viroporins in the Viral Life Cycle and Their Regulation of Host Cell Responses. Frontiers in Immunology, 13. 10.3389/fimmu.2022.890549

47. Zhang, M., Zeng, C. Q.-Y., Morris, A. P., & Estes, M. K. (2000). A Functional NSP4 Enterotoxin Peptide Secreted from Rotavirus-Infected Cells. Journal of Virology, 74(24), 11663–11670. 10.1128/JVI.74.24.11663-11670.2000

